# Auditory Countermeasures for Sleep Inertia: An Ecological Study Examining the Influence of Melody and Rhythm

**DOI:** 10.1101/2020.03.03.974667

**Authors:** Stuart J. McFarlane, Jair E. Garcia, Darrin S. Verhagen, Adrian G. Dyer

**Author notes:** (SJM).

## Abstract

Sleep inertia is the potentially harmful decline in cognition that occurs upon and following awakening. Sound has been shown to counteract the negative symptoms of sleep inertia, with a recent study revealing that an alarm perceived as melodic by participants displayed a significant relationship to reports of reductions in perceived sleep inertia. This current research builds on these findings by specifically testing the effect melodic and rhythmic stimuli exhibit on sleep inertia for subjects awakening in their habitual environments. Two test Groups (A & B; *N* = 10 equally) completed an online psychomotor experiment and questionnaire in two separate test sessions immediately following awakening from nocturnal sleep epochs. Both groups responded to a Control stimulus in the first session, while in the second session, Group A experienced a Melodic treatment, and Group B the Rhythmic. The results show that the melodic treatment significantly decreased attentional Lapses, False Starts and had a significantly improved PVT Performance Score than the Control. There was no significant result for Reaction Time or Response Speed. Additionally, no significant difference was observed for all PVT metrics between the Control – Rhythmic conditions. The results support melodies potential to counteract symptoms of sleep inertia by the observed increase in participant vigilance following waking. Specifically, a melodically rhythmic contour is highlighted as a significant musical treatment noteworthy of consideration when designing alarm compositions for the reduction of sleep inertia. As auditory assisted awakening is a common within modern society, improvements in alarm sound design may have advantages in domestic and commercial settings.

## Introduction

Sleep inertia (*SI*) is a transitional sleep-wake phenomenon defined by a reduction in human performance upon, and post-awakening (1–3). Originally examined through the testing of human performance decrements upon sudden awakening (4–6), subsequent research conducted by Lubin et al (7) described the phenomena as ‘nap inertia’, of which provides the basis for the current terminology referred to in practice today.

The adverse features of *SI* have been shown to protract for approximately 0 - 30 minutes post-awakening, however durations spaning up to 4 hours have also been reported (3, 8–13). Studies have shown *SI* to impair several dimensions of cognitive performance, including reaction time (RT) (14–16), and decision making (17, 18). In a real world context, Wertz et al (8) suggests that the resulting decline in performance may be on par, or more pronounced than being legally intoxicated, and/or a night of complete sleep deprivation.

Deficits in human performance post-awakening may have serious costs for personnel working in high risk positions; particularly in the areas of health, emergency response, and vehicle control; where human error through lack of cognition may prove detrimental (19, 20).

A number of factors have been researched in an attempt to further understand and manage *SI*, of which include awakening countermeasures; also referred to as reactive countermeasures (21), environmental factors (3), and experimental manipulation (1). Awakening countermeasures have been researched in several contexts and themes, which may be considered as extensions or augmentations of ubiquitous human behaviours and routines. These include; caffeine (14, 22–25), light (16, 24, 26–28), temperature (29, 30), post-awakening routines (24, 31), sound (32–34), and stress (35).

With respect to this current research investigation, three previous studies have been identified which examine how audio may impact *SI* post-awakening. Tassi et al (32) concluded that pink noise (75 dB) can reduce *SI* when deployed as an intense waking alarm, while Hayashi (33) detected that excitatory music, particularly high-preference popular music (60 dB) as specified by participants has the potential to reduce the impact of *SI* after a short nap. Through an ecologically valid approach McFarlane et al (34) revealed that participant alarm sounds perceived as melodic showed a significant relationship to reductions in perceived sleep inertia, as compared to ‘neither unmelodic nor melodic’ counterparts. Additionally, it was shown that a melodic alarm sound is perceived to be more rhythmic than a neutral interpretation (34). Taken together, these three studies discussed above demonstrate that sound and music are plausible awakening countermeasures for *SI*, and with additional research we may establish a refined understanding of the auditory aesthetics and musical mechanisms required for the best practice design of such stimuli.

Noise, sound and music have been shown to enhance arousal and improve task performance in alert humans. For example, loud white noise (85 – 100 dB) may improve simple addition (36), and attentional selectivity (37) when compared to quieter signals (52 dB and 65 dB respectively). Visual vigilance discrimination has been shown to be enhanced relative to a musical counterpart (38). More broadly, Poulton (39) indicates that noise can enhance performance, particularly for tasks requiring speed or vigilance. With respect to sound and ‘noises’, Asutay and Västfjäll (40) suggest that environmental sound (e.g. boiling water, fingernails on a blackboard, toilet brush) facilitates improved visual attention when compared to quiet conditions.

Instrumental music has also been observed to be beneficial for the improvement of sustained attention. Davies, Lang and Shackleton (41) analysed the effectiveness of noise and instrumental music on visual attention during two visual vigilance task conditions: difficult and easy. In this analysis, music was shown to significantly prevent detection latencies in the difficult task condition. All stimuli deployed had an approximate loudness of 75 dB, with the instrumental music conditions including a solo guitar performance by Laurindo Almeida, and an orchestration by Don Ellis, however, the specific compositions were not reported.

‘Rock music’ has been reported to improve signal detection compared to ‘instrumental music’ (75 dB respectively) (42), and task performance (43). For example, while working, three subject groups where exposed to fast-paced (instrumental ‘rock’ music, ~140 beats per minute [BPM]), slow-paced (instrumental ‘heartbeat’ music, 60 BPM), and a no music control condition. Participants undertook two associated activities; (i) looking up and recording closing stock prices during October-December 1987, and; (ii) calculating percentage changes for each week during this period. Mayfield and Moss (43) deduced that task performance was higher in the rock music condition than in both the ‘heartbeat’ music and no music conditions, though the authors note that participants reported a significantly increased subjective distraction rating in the rock music condition. In similar reporting to Davies, Lang and Shackleton (41), the authors present minimal musical detail regarding the test stimuli the participants are experiencing.

Melodically rhythmic instrumental music is reported to have increased human task performance (44, 45). For instance, while recording Event-Related Potential (ERP) brain activity, Riby (44) tested seventeen participants performing a visual odd-ball task (a rare target stimulus, a rare novel stimulus, and a frequent nontarget stimulus) when listening to each of Vivaldi’s Four Seasons concertos’; Spring, Summer, Autumn, and Winter, relative to a silent control. It was shown that the ‘Spring’ concerto enhanced subject’s mental alertness, attention and memory when compared to the silent control and the ‘Summer’, ‘Autumn’, and ‘Winter’ selections (44). The authors suggest that ‘Spring’s’ improvement in task performance may be attributed to the major mode of the piece, and the faster tempo of its first movement (44), which they in turn hypothesized enhanced the perceived vibrance and positive emotion of the participants; leading to arousal. ‘Spring’ is performed in E Major at three tempos throughout the concerto (Allegro [120 BPM - 156 BPM]; Largo [40 BPM – 60 BPM]; Allegro [120 BPM - 156 BPM]) (46).

From an operational and end-user perspective, real-world circumstances may benefit from sound stimuli targeted at the reduction of *SI*, including occupational settings where audio is employed to activate employees immediately following awakening, and improved day to day awakening alarm tones, of which are common with in society and our auditory ecology (47). In the current study we examine the influence saliently melodic and rhythmic alarm tone treatments have on *SI* immediately post-awakening in ecological conditions comparative to a non-melodic control. Through this investigation we aim build on McFarlane et al’s (34) initial findings to further understand if and how melody may assist in counteracting *SI*, and the relationship rhythm may have to *SI* reduction when tested in isolation. Within the broader context of auditory assisted awakening, our goal is to provide evidence that may be expanded upon and referenced in the future development and testing of sound stimuli to counteract *SI* in natural, auditory complex surroundings.

## Materials and Methods

### Ethics Statement

All research methods, participant numbers (*N* = 20) considered appropriate for the study, and data collection were approved by the Royal Melbourne Institute of Technology University’s s (RMIT) College of Human Ethics Advisory Network (CHEAN) (ref: CHEAN B 21753-10/18). Respondents provided their specific consent to participate by completing the online study. This was stipulated to the subjects in the ‘Invitation to participate’ email distributed during the recruitment period, and reiterated prior to undertaking the online test. The study was launched during May 2019 and concluded in November 2019.

### Participants

Subjects were invited to participate through RMIT’s School of Media and Communications staff, student and membership networks, printed posters located throughout the RMIT University Melbourne city campus, and through the researches social networking communities. Individuals interested in volunteering for the study contacted the lead researcher directly via email. The volunteers were then supplied the study’s ‘Invitation to participate’ form. The contents included an introduction, title and overview of the research, who is conducting the study, participants’ rights and responsibilities, instructions for how to undertake the test, and contact information for any further enquiries regarding the test. Ideal participants were required to be 18 years and above, healthy with good hearing, have a consistent sleeping pattern and access to a smart phone, computer, tablet or laptop, and a secure internet connection. Participants where not renumerated for their service. All eligible participants were encouraged to undertake the study without bias towards music preference or training.

The recruited participants gender classification was obtained through a Male, Female, and X (Intermediate / Intersex / Unspecified) question in reference to the Australian Government guidelines on recognition of sex and gender (48). Further, an option of non-disclosure was included to accommodate any participant willing to volunteer, yet reticent in recording any gender classification (i.e. Prefer not to disclose). We have refrained from reporting or supplying the gender demographics for each individual analysis of the participants as open access data to support data minimization. This research reporting strategy is consistent with the General Data Protection Regulation (GDPR) (49).

### Data Collection

The reported data was captured digitally via the use of the online software system Gorilla (50), where the questionnaire and experiment is contained, managed, and remotely accessed. Gorilla is software specifically produced for the undertaking of online questionnaires and experiments enabling researchers to design and implement their studies for ethically compliant distribution and data collection. The data obtained by Gorilla is securely stored and available for download and analysis by researchers. Gorilla is fully compliant with GDPR (50), and is developed with reference to The British Psychological Society (51), and National Institute for Health Research (52) standards.

### Test stimulus – Design and Description

All stimuli are in the key of C, have a meter of 4:4, a tempo of 105 BPM, and comprise a monophonic (Control) and polyphonic (Melody, Rhythm) texture. The arrangements where designed as a two-bar motif and repetitively looped to a total duration of 108 seconds for Android or PC users, and 2 bars for Apple users due to the specific audio play-back features of each platform. All compositions have been produced in the audio production software package Cakewalk (53) and employ the TTS – 1 soft synth to trigger the Vibraphone W and Woodblock timbral sample sets from the sound library provided. The final compositions are digitally limited and compressed to produce clear and balanced audio, which are exported as MP3/4 files. Caution was applied during the design phase to the auditory performance of the stimuli when relayed through various multimedia devices (e.g. mobile phones, laptops and tablets). Lower frequencies do not perform as effectively as higher tones due to the limited frequency range these device types can produce (54). All stimuli were iteratively prototyped through extensive field testing during the design development period (2018 - 2019).

The objective for the auditory design of the Control, Melodic and Rhythmic test stimulus was to produce a set of three recognizable, yet original complementary compositions, that when qualitatively compared, are differentiated and easily interpreted by their individual musical attributes (i.e. Control [Neither overtly rhythmic nor melodic]; Melodic, and Rhythmic). In this way we established a framework where the stimulus designed, in conjunction with the experimental study design, enables the elemental analysis of each stimulus and their effect on *SI*.

To achieve this, we first produced the Control as a metronomic pulse that can be sounded independently and perform as the ‘heartbeat’ to both the Melodic and Rhythmic stimulus. The Melodic and Rhythmic stimuli are layered upon the Control and strategically composed to accentuate their elemental musical aesthetics by means of timbre and contour. See Table 1 for the design overview of the stimulus set.

**Table 1.** Stimulus design overview.

| TEST STIMULI DESIGN OVERVIEW |  |  |
| --- | --- | --- |
| CONTROL | MELODY | RHYTHM |
| Metronomic contour<br>(Woodblock, Vibraphone) | Metronomic contour<br>(Woodblock, Vibraphone)<br>+<br>Melodic contour<br>(Vibraphone) | Metronomic contour<br>(Woodblock, Vibraphone)<br>+<br>Rhythmic contour<br>(Woodblock) |

The key of C was deemed appropriate for the three stimuli in this study’s context considering its extensive application in popular music, and universal familiarity (55, 56). Similarly, the 4:4 meter, also known as ‘Common time’ (57), was selected as it is the most frequently employed and recognisable time signature in Western music today (56, 58, 59).

A tempo of 105 BPM was established as an appropriate pace for the function of the stimuli we sought to achieve, which is to successfully enable awakening, yet not to be overtly alarming, salient or fast, nor slow or calming. Residing in the range of the classical andante tempi (76 BPM - 108 BPM) (60), the preferred perceptual tempo (PPT) (also identified as preferred tempo and indifference interval) (61, 62) of 100 BPM as proposed by Fraisse (62) and marginally slower than Moelants (61) finding of 120 BPM. A tempo of 105 BPM may be described as an approximate ‘mid-range’ with respect to the human tempo registration range (existence region) of 40 BPM - 300 BPM (63). This method has been chosen to allow for the targeted within subject comparisons between the respective musical treatments without the influence of tempo (slow paced – fast paced) on arousal.

The Control stimulus comprises two timbres (Woodblock, Vibraphone) that are sounded on every 1^st^ and 3^rd^ beat of each bar. The Woodblock timbre is the percussive element of the score and is denoted the single note D6 with respect to an 88 key virtual piano (100% velocity: 240 ms duration). The tonal element of the Control layers the Vibraphone sample as a single C6 note in relationship to an 88 key virtual piano (100% velocity: 240 ms duration) over the D6 percussive notes. When played, the Control produces a metronomic and inexpressive pulse.

The Melodic stimuli retains the D6 (100% velocity; 240 ms) percussive timbre and arrangement of the Control, yet strategically increases the vertical tonal contour, and horizontal rhythmic contour of the Vibraphone W notes within the composition. By introducing musical notes C7, A6, G6, E7 and E6 to the Control, reducing inter-onset intervals (IOI’s) between notes, and enhancing the tonal contour, the resulting passage is designed to generate a dramatic rise in perceived melodicity when compared to the Control. The dynamic aesthetic of this composition employs variations in note velocity (85% - 100%), and duration (174 ms – 340 ms).

The Rhythmic stimuli retains the horizontal rhythmic contour and dynamic features of the Melodic composition, however, restricts the vertical tonal contour and timbre of the score to D6 and Woodblock respectively. In so doing, the composition may be interpreted as the rhythmic counterpart to both the Control and Melodic scores through the increased rhythmic contour (comparative to the Control) and salient percussive timbre (relative to the Melodic stimuli). See Fig 1. for the stimulus musical notation.

**Fig 1.**
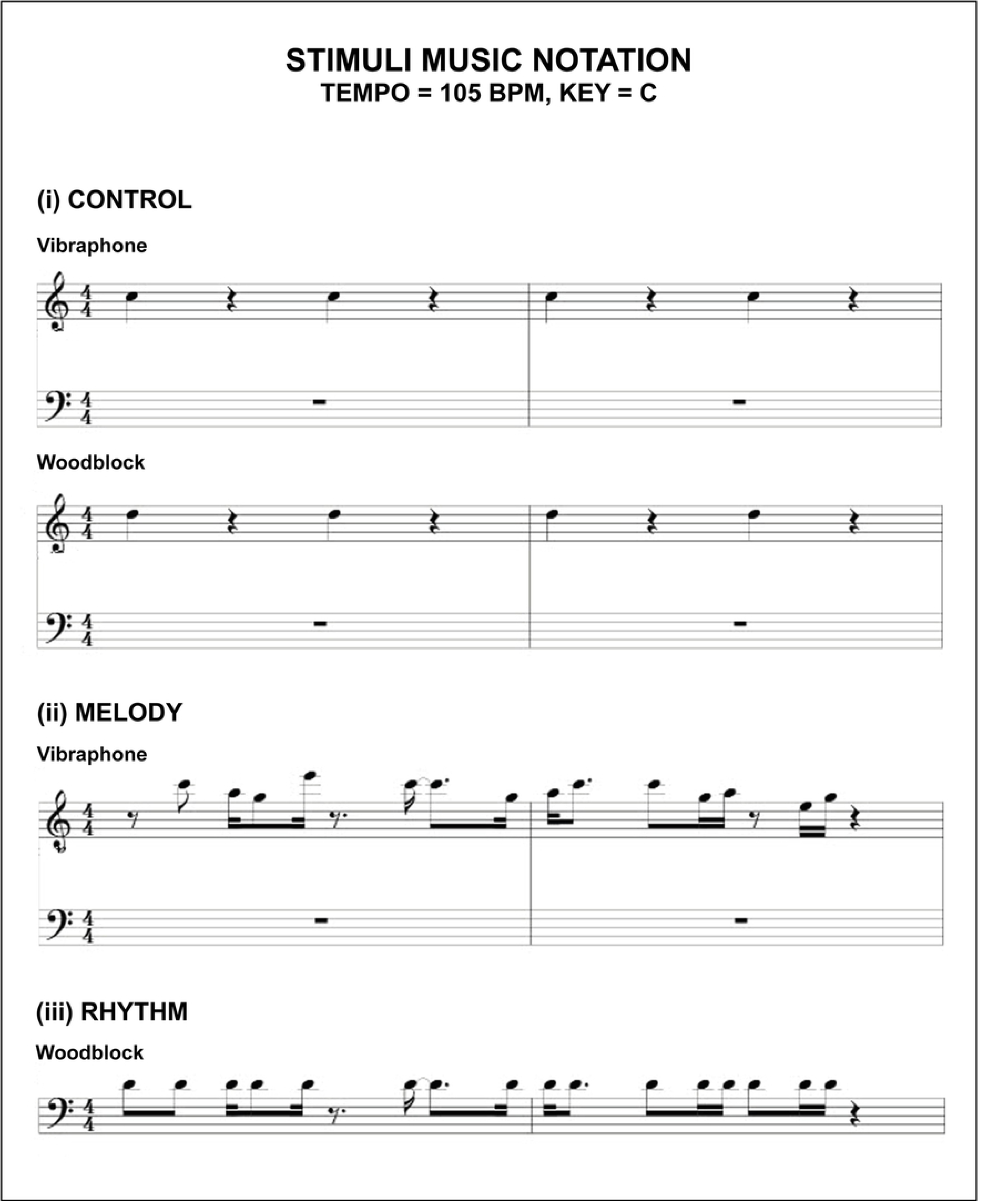
Musical notation for the Control, Melodic, and Rhythmic stimuli.

### Study Design

The study comprised of two Groups (A & B) that were required to undertake two test sessions (Session 1 & Session 2) conducted each week on a Tuesday and Wednesday morning respectively during the data gathering window. All participants were required to complete each test immediately after waking from an assigned stimulus that was supplied by the researchers as a replacement to their usual alarm sound. The waking stimulus was deployed on the participants preferred electronic device (desktop, laptop, tablet, or smart phone). Participants completed each test in their chosen location and at their typical time of waking. This method was selected to maximise the natural contextual environment in which subjects use auditory alarms for awakening in their daily routine, ensuring the ecological validity of the findings. Each test session requires approximately 5 - 10 minutes to complete, being designed to collect high value data whilst minimising disruption to participants.

The Participants (*N* = 20) were pseudo-randomly allocated and equally divided into two groups (Group A, *N* = 10; Group B, *N* = 10). Session 1 (Tuesday) required both groups to complete the study after awakening to the Control stimulus. Session 2 (Wednesday) required Group A to complete the test following awakening to the melodic stimuli, and Group B the rhythmic stimuli. Table 2 illustrates the study protocol.

**Table 2.** Study protocol diagram.

| EXPERIMENTAL TEST PROTOCOL |  |  |  |
| --- | --- | --- | --- |
| SUBJECT GROUPS | PRE-TEST PREPORATION | SESSION 1. | SESSION 2. |
| GROUP A | Test hyperlink<br>Instruction document<br>Stimuli audio files x 2 | CONTROL STIMULUS | MELODY STIMULUS |
| GROUP B | Test hyperlink<br>Instruction document<br>Stimuli audio files x 2 | CONTROL STIMULUS | RHYTHM STUMULUS |

At a minimum of twenty-four hours prior to commencing the study each test group were supplied email the test hyperlink via, instruction document (PDF), and the test stimuli audio files. The hyperlink allowed each participant to access the test on the first day and resume the test the following morning. The design of the online study included timing nodes to safeguard against any participant attempting to access the study prior to (or between) each test date. The instruction document contained the pre-test preparation and the test procedure.

The pre-test preparation consisted of six steps for each participant to follow. These included: Setting up the sounds on your device (Step 1), Setting up the sounds as your two alarms (Step 2), Setting the alarm volume (Step 3), Testing the alarm sounds (Step 4), Email link to the study (Step 5), and Test preparation (Step 6). Steps 1 – 4 instruct each participant to first download the test stimuli onto their device, set the files as a separate alarm tones (each stimuli file is labelled corresponding to Tuesday or Wednesday), define the volume and disable the ‘rising volume’ setting if applicable, test both stimuli for correct functionality, and familiarize themselves with each stimulus. Step 5 informs the participants that the hyperlink will direct them to the study and to check their email accounts as reminder emails will be issued prior to the second days test. Step 6 recommends that each participant has the relevant email open the night prior in preparation to activate the study link.

The test procedure information commences by encouraging each participant to familiarize themselves with the protocol prior to undertaking the test, and is followed by the process for each test session. The procedure for each session contains three steps for each participant to follow under the themes; Upon Waking (Step 1); Beginning the Test (Step 2); End of the Session (Step 3).

The test battery for each study included an adapted brief psychomotor vigilance task (PVT, 3 min [Item 1]) (64), the Karolinska Sleepiness Scale (KSS, Item 2) (65–67), and two custom designed Likert scales (Sleep Duration [Item 3]; Sleep Quality [Item 4]). Session 1 also recorded Demographic Information (Gender [Item 5]; Age range [Item 6]; Hours typically slept [Item 7]). Please refer to the supporting material (S1 Table) for a transcript of the questionnaire for each test session. All responses were forced.

The PVT-B (68) is a validated 3-minute variation of the popular 10-minute PVT developed by Dinges and Powell (69) which records participant reaction time (RT) to random interval stimuli. Interpretation of this data is extrapolated into several performance metrics (i.e. mean RT, Lapses and False Starts) as measures of behavioural alertness. Several research experiments have incorporated the PVT-B as an objective measure of a subject’s vigilance (70, 71). Benefits for using the PVT-B in this study’s context are that it can be undertaken remotely online and is intuitive for participants to perform (72). The PVT-B is particularly suited to enquiries where the 10-min PVT is considered overtly time consuming (64). Our adapted PVT-B was produced in the Gorilla task builder software (50) with respect to Basner and Dinges (68) and Basner et al (64). The test requires each participant to either click a mouse controller, depress a keyboard key, or press a button icon on a screen (dependant on which device the participant nominates) immediately as a visual stimuli transitions from one assigned colour to another. In our design the subjects are instructed to respond as quickly as possible when a circular orange stimulus turns red. The interstimulus interval (ISI) between each coloured stimulus was randomized and varied between 1 and 4 seconds as specified by Basner et al (64). A timeout condition was included (≥ 1000 ms) to further ensure test duration is retained to a minimum while retaining responsiveness. During analysis, timeouts are interpreted as Lapses with a 1000 ms duration. One of three statements are displayed following each response as a fixation substitute and to inform the participant of their continual performance. These are: (i) Too Quick! (False Start); (ii) Great Work! (Correct response); Too Slow! (Timeout). Each statement extends for 1000 ms and is deducted from the total (ISI) as previously described.

The KSS (65) is a subjective 9-point bipolar Likert scale measure of sleepiness that exists in two versions, A and B (73). The original KSS A labelled the odd scales only (1 = Extremely alert, 3 = Alert, 5 = Neither alert nor sleepy, 7 = Sleepy but no effort to keep awake, and 9 = Very sleepy and a great effort to keep awake, fighting sleep) while the KSS B (66) subsequently completed the even labels (2 = Very alert, 4 = Rather alert, 6 = Some signs of sleepiness, and 8 = Sleepy, some effort to keep awake). These two versions have been verified to be similar (73) and results comparable. In this study we have implemented version B. The instructions request the participant to indicate their ‘level of sleepiness during the 5 minutes before this rating by selecting the appropriate description’. We interpret the response of this rating to reflect the perceived sleepiness of participants upon waking prior to the commencement of the PVT.

The custom designed subjective measure of sleep duration (Item 3) is a self-report unipolar 14-point Likert scale. This design requests each participant to rank their sleep duration as “accurately as possible” from the fourteen options supplied. Each sleep time category is measured in increments of 0.5 hours between either end categories of the scale (0 – 3 hours and 9+ hours). For example, 0 – 3, 3 – 3.5, 3.5 – 4, 4 – 4.5 etc.

To record each participants’ subjective sleep quality, we developed a self-report 5-point unipolar Likert scale. The scales options for selection contain, Very Poor, Poor, Average, Good, and Very Good. The decision to design this single item scale as opposed to utilizing the established Sleep Quality Scale (SQS, 5 - 10 min completion time) (74) or the Pittsburgh Sleep Quality Index (PSQI, 1-month reporting duration) (75), was to reduce potential time constraints of each participant. As the test is undertaken prior to each subject attending their employment obligations (if applicable), limiting the test duration was a factor in the design of this study. To gather demographic data of the participants typical hours slept each night we included a 5-point unipolar Likert scale with the following options in hours, 0 - 3, 3 - 5, 5 - 7, 7 - 9, and 9+.

### Statistical Analysis

With reference to Basner and Dinges (68) the PVT performance metrics recorded in this study are the mean RT, mean 1/RT (reciprocal response time or response speed calculated by dividing RT (ms) by 1000, then reciprocally transformed) (76), the number of Lapses (≥ 500 ms), the number of False Starts (RT ≤ 100 ms and responses prior to the red stimuli), and Performance Score (i.e. 1 minus the number of Lapses and False Starts divided by the number of valid stimuli including False Starts) (68).

A planned sequence of five paired sample t-test’s (77) were completed to examine each vigilance metric between conditions for the respective groups (Group A: Control – Melody; Group B: Control – Rhythm). Cohen’s *d* is the effect size employed for the analysis of the paired t-test’s and is calculated by the mean of within-subject differences, divided by the standard deviation of the within subject differences (78). The Shapiro-Wilk test was applied to asses normal distribution of the data (79). Data sets which reject the normality hypothesis were analysed using the non-parametric Wilcoxon signed rank test (80).

The Wilcoxon signed rank test was also employed to analyse the median rank within subject differences for the subjective measures (KSS, Sleep Duration, Sleep Quality) in each individual test group. The effect size selected for the analysis of each Wilcoxon signed rank test was calculated from the *z*-value reported by the test, divided by the square root of the number of observations recorded (i.e. *N* = 20) (81). Additionally, percentage comparisons were undertaken for each subjective measure as an aggregate of both test sessions for each test group. An α = 0.05 was considered statistically significant for the analysis. Raw data was tabulated for analyses in Microsoft Excel (82), then imported to SPSS 26 (83) for statistical analysis.

## Results

### Test Group A: Melody Stimulus

Participants were aged between 18 and 49 (18 – 29 [n = 5], 30 – 39 [n = 2], 40 – 49 [n = 3]), consisted of Males and Females (40% Female), with 40% of participants reporting consistent sleep epochs of 5 - 7 hours per night, and 50% 7+ hours respectively (3 – 5 hours [n = 1], 5 – 7 hours [n = 4], 7 – 9 hours [n = 4], 9+ hours [n = 1]).

### Group A PVT Metrics

**Fig 2.**
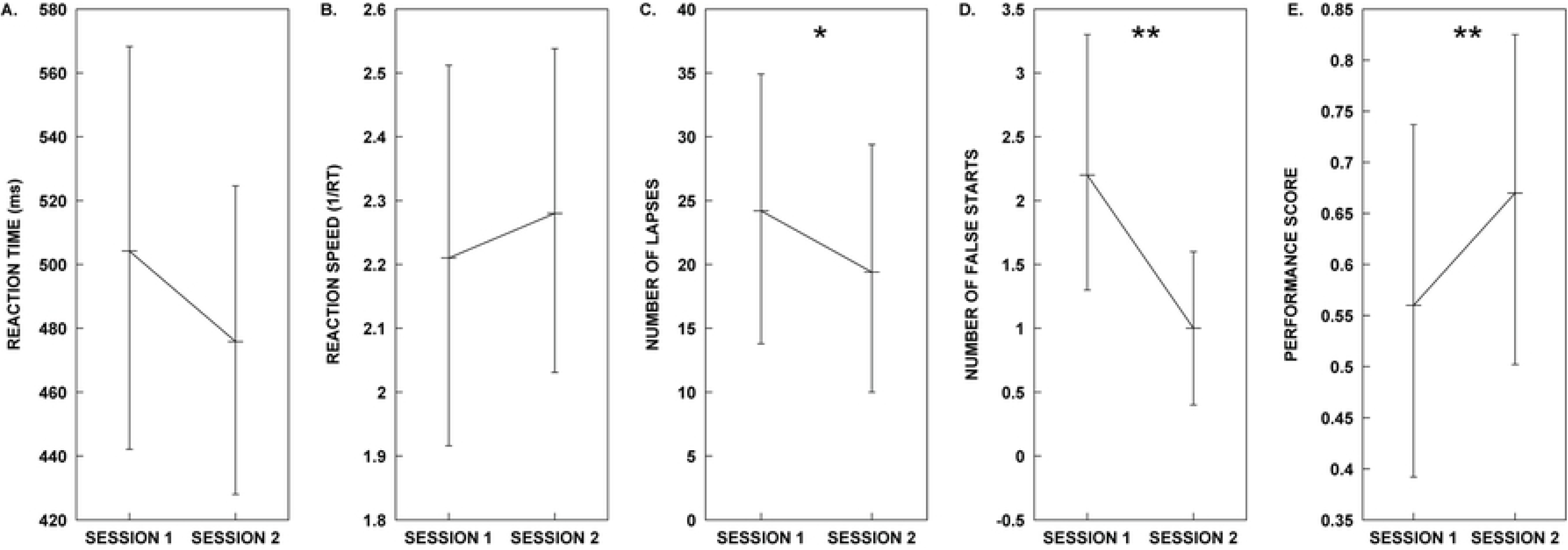
Test Group A plots for PVT metrics in each test session.

Results for the five different PVT metrics obtained from participants in Group A during sessions 1 and 2. (A) mean Reaction Time, (B) mean Reaction Speed, (C) mean number of Lapses, (D) mean number of False Starts and (E) mean Performance Score. We identified significant differences in performance at alpha = 0.05 for Lapses, False Starts and Performance. P-values less than 0.05 are represented by (*). P-values less than 0.01 are represented by (**). All error bars represent 95% confidence intervals. Refer to the Results section for details on the outcome of the statistical analyses.

The planned sequence of five paired-sample t-tests were performed to analyse the PVT metrics (mean RT, mean 1/RT, Lapses, False Starts, and Performance Score) within subjects for the Control (Session 1) and Melody (Session 2) conditions. Our results show that there is no significant difference with a medium effect (≥ 0.5) (78) in the mean RT for the Control and Melody conditions, nor a significant difference between conditions with a small effect (≥ 0.2) (78) in mean inverse reaction time (mean 1/RT). The analysis does reveal a significant difference with a large effect (≥ 0.8) (78) between the mean Lapses, mean False Starts, and the PVT Performance Score results for the Control and Melody treatments. Specifically, the Melody treatment resulted in superior performance considering Lapses, False Starts and the PVT Performance Score. See Table 3.

**Table 3.** PVT metrics of the paired sample t-test results for Test Group A (Melody Stimulus).

| MEASURE | SESSION 1. CONTROL | SESSION 2. MELODY | TEST STATISTICS |
| --- | --- | --- | --- |
| Mean RT | $\mu = 504.21, SD = 107.02$ | $\mu = 475.84, SD = 84.43$ | $t(9) = 1.44, p = 0.184, d = 0.455$ |
| Mean 1/RT | $\mu = 2.21, SD = 0.50$ | $\mu = 2.28, SD = 0.43$ | $t(9) = -0.681, p = 0.513, d = -0.215$ |
| Lapses | $\mu = 24.20, SD = 18.29$ | $\mu = 19.40, SD = 17.10$ | $t(9) = 2.94, p = 0.016, d = 0.930^*$ |
| False Starts | $\mu = 2.20, SD = 1.69$ | $\mu = 1.00, SD = 1.05$ | $t(9) = 4.811, p = 0.001, d = 1.521^{**}$ |
| Performance Score | $\mu = 0.56, SD = 0.30$ | $\mu = 0.66, SD = 0.28$ | $t(9) = -3.54, p = 0.006, d = -1.121^{**}$ |

### Group A Subjective Measures

**Fig 3.**
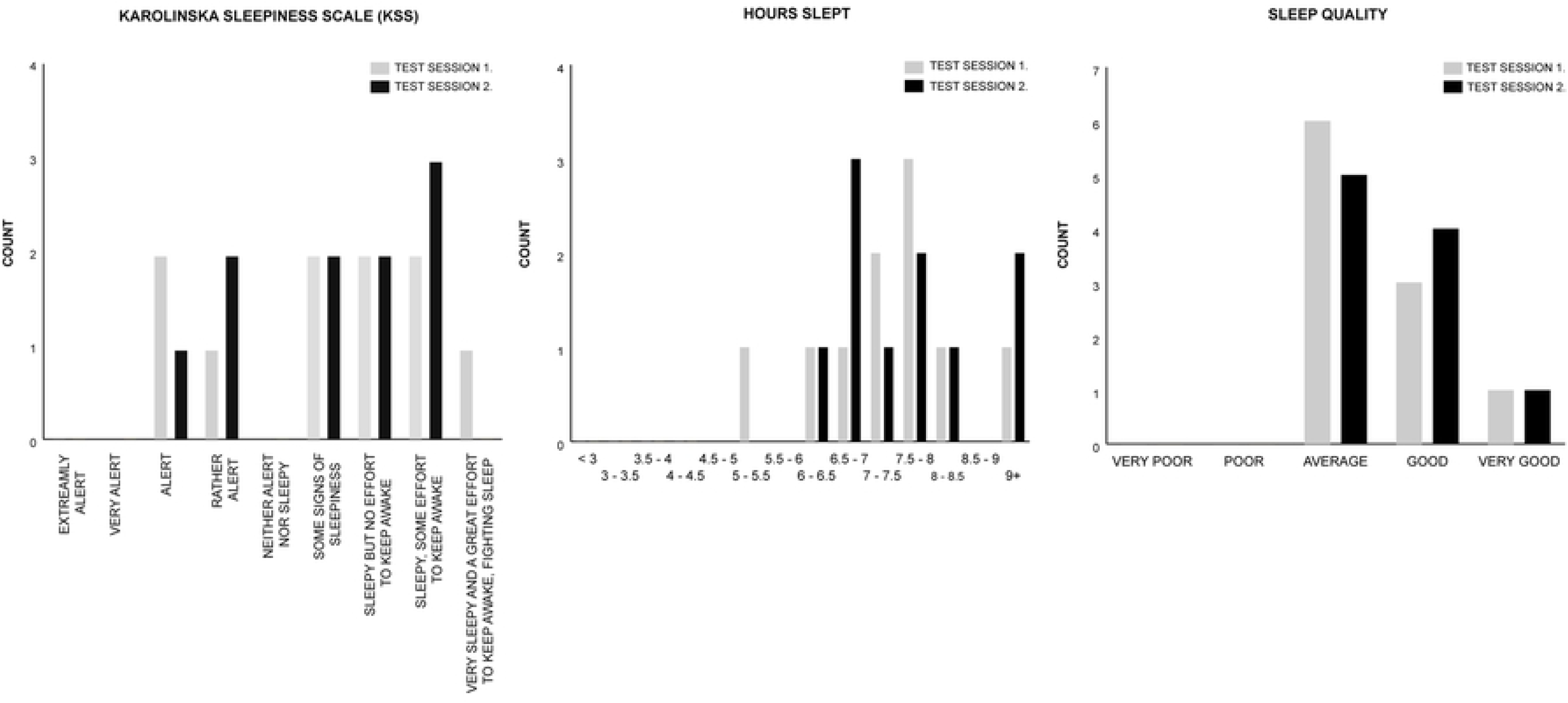
Test Group A histogram counts for the KSS, Hours Slept, and Sleep Quality measures.

The Wilcoxon signed rank test was utilized to individually analyse the median difference of the KSS, Hours Slept, and Sleep Quality measures within subjects for the Control (Session 1) and Rhythm (Session 2) conditions of Test Group A. For all measures the median Session 2 test ranks are not significantly different than the Session 1 test ranks, indicating that in both test sessions participants subjective sleep attributes are analogous. See Table 4.

**Table 4.** Wilcoxon signed rank test results for subjective measures of Test Group A.

| MEASURE | SESSION 1. CONTROL | SESSION 2. MELODY | TEST STATISTICS |
| --- | --- | --- | --- |
| KSS | $M = 6.50$ | $M = 6.50$ | $z = 0.000, p = 1.000, r = 0.000$ |
| HOURS SLEPT | $M = 10.50$ | $M = 10.50$ | $z = -1.186, p = 0.236, r = -0.265$ |
| SLEEP QUALITY | $M = 3.00$ | $M = 3.50$ | $z = -0.378, p = 0.705, r = -0.085$ |

Mean percentages across the two test sessions for each subjective measure show that 70% of participants reported ratings of ‘sleepiness’ (Some signs of sleepiness [20%]; Sleepy but no effort to keep awake [20%]; Sleepy, some effort to keep awake [25%]; Very sleepy and a great effort to keep awake, fighting sleep [5%]), while 30% reported to be ‘Rather Alert’ (15%) to Alert (15%). The mean hours slept prior to each test session reveal that 65% of participants slept 7+ hours (6+ hours [95%]); and 100% reported ratings from ‘Average’ to ‘Very good’ sleep quality (Average [55%]; Good [35%]; Very Good [10%]).

### Test Group B: Rhythm Stimulus

Participants were aged between 18 and 49 (18 – 29 [n = 4], 30 – 39 [n = 5], 40 – 49 [n = 1]), consisted of Males and Females (50% Female), and reported a typical night’s sleep ranging from 5 – 9 hours (5 – 7 hours [n = 5; 50%], 7 – 9 hours [n = 5, 50%]).

### Group B PVT Metrics

**Fig 4.**
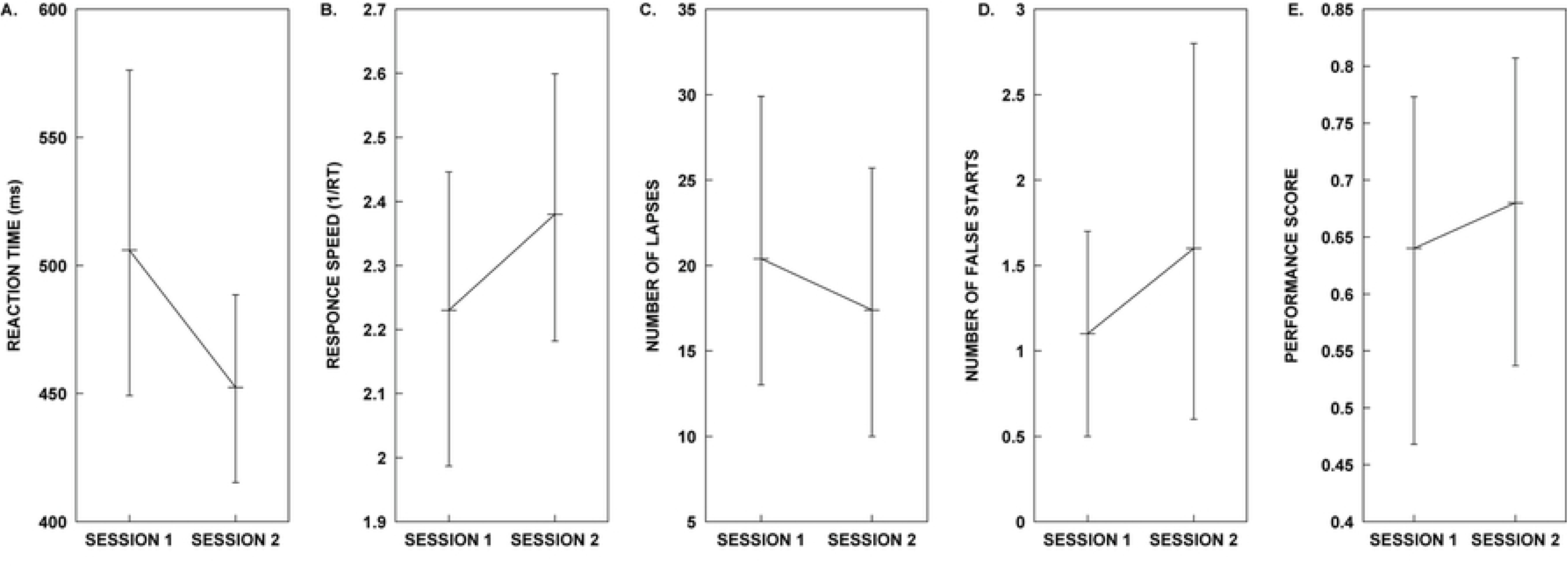
Test Group B plots for PVT metrics in each test session.

Results for the five different PVT metrics obtained from participants in Group A during sessions 1 and 2. (A) mean Reaction Time, (B) mean Reaction Speed, (C) mean number of Lapses, (D) mean number of False Starts and (E) mean Performance Score. All error bars represent 95% confidence intervals. Refer to the Results section for details on the outcome of the statistical analyses.

Consistent with Test Group A’s analysis, planned paired-sample t-tests were performed to analyse the PVT metrics within subjects for the Control (Session 1) and Rhythm (Session 2) conditions. Our results show that there is no significant difference in the mean RT, mean 1/RT, Lapses, nor Performance Score for the Control and Rhythm conditions. The Wilcoxon signed ranks test for False Starts indicate that the median Session 2 test ranks are not significantly different than Session 1 test ranks (Table 5).

**Table 5.** PVT metrics analysis results for Test Group B (Rhythm Stimulus).

| MEASURE | SESSION 1. CONTROL | SESSION 2. RHYTHM | TEST STATISTICS |
| --- | --- | --- | --- |
| Mean RT | $\mu = 506.01, SD = 109.68$ | $\mu = 452.42, SD = 64.22$ | $t(9) = 1.405, p = 0.193, d = 0.444$ |
| Mean 1/RT | $\mu = 2.23, SD = 0.40$ | $\mu = 2.38, SD = 0.35$ | $t(9) = -1.058, p = 0.318, d = -0.335$ |
| Lapse | $\mu = 20.40, SD = 14.92$ | $\mu = 17.40, SD = 13.45$ | $t(9) = 0.530, p = 0.609, d = 0.168$ |
| False Starts | $M = 1.00$ | $M = 1.00$ | $z = -1.30, p = 0.194, r = -0.291$ |
| Performance Score | $\mu = 0.64, SD = 0.25$ | $\mu = 0.68, SD = 0.23$ | $t(9) = -0.421, p = 0.683, d = -0.133$ |

### Group B Subjective Measures

**Fig 5.**
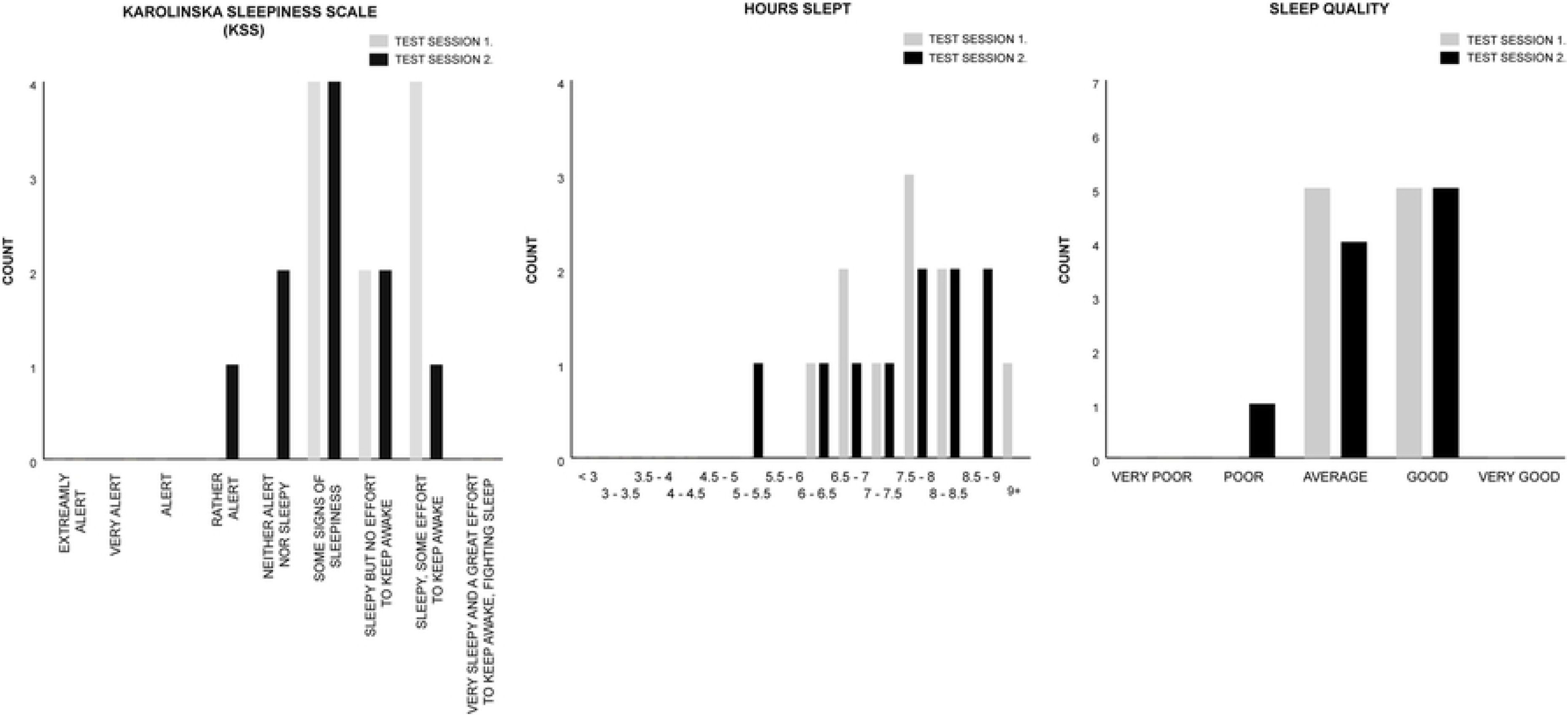
Test Group B histogram counts for the KSS, Hours Slept, and Sleep Quality measures.

The Wilcoxon signed rank test was utilized to individually analyse the median difference of the KSS, Hours Slept, and Sleep Quality measures within subjects for the Control (Session 1) and Rhythm (Session 2) conditions of Test Group B. For all measures the median Session 2 test ranks are not significantly different than the Session 1 test ranks, indicating that in both test sessions participants subjective sleep attributes are comparable (See Table 6).

**Table 6.** Wilcoxon signed rank test results for subjective measures of Test Group B.

| MEASURE | SESSION 1. CONTROL | SESSION 2. RHYTHM | TEST STATISTICS |
| --- | --- | --- | --- |
| KSS | $M = 7.00$ | $M = 6.00$ | $z = -1.699, p = 0.089, r = -0.380$ |
| HOURS SLEPT | $M = 11.00$ | $M = 11.00$ | $z = -0.431, p = 0.666, r = -0.096$ |
| SLEEP QUALITY | $M = 3.50$ | $M = 3.50$ | $z = -0.577, p = 0.564, r = -0.129$ |

Group B’s mean percentages across both test sessions for each sleep measure show that 85% of all participants reported ratings of ‘sleepiness’ (Some signs of sleepiness [40%]; Sleepy but no effort to keep awake [20%]; Sleepy, some effort to keep awake [25%]). One hundred percent (100%) of Session 2 participants reported a minimum rank of ‘sleepiness’. Thirty percent of Session 1 participants reported not to be sleepy (‘Neither Alert nor Sleepy’ [20%]; Rather Alert [10%]). Seventy percent (70%) of all Group B participants slept 7+ hours (6+ hours [95%]); and 95% reported ratings from ‘Average’ to ‘Good’ sleep quality (Average [45%]; Good [50%]).

## Discussion

The objective of this study was to assess the effect musically melodic and rhythmic auditory alarm tone treatments exhibit on symptoms of *SI* following awakening within ecological conditions. To date this research represents the first experiment to test and report reproducible alarm tones with strategically composed melodic and rhythmic contours in the context of *SI*. The results obtained present key insights that may be utilized to extend our understanding for how alarm tone design may assist to counteract *SI*, which may prove beneficial in scenarios where sustained attention is vital immediately upon waking. Examples would include land based, aeronautical, and nautical transportation; critical monitoring tasks; or common day to day activities like driving, or riding a bike post-awakening.

The principal results of this study reveal two key findings: (i) A saliently Melodic alarm tone, incorporating a musically neutral Control test stimulus within its design, enhanced vigilance post-awakening when compared to the Control stimulus in isolation. (ii) A Rhythmic alarm tone comprising of the identical rhythmic contour as the Melodic stimuli with the Control imbedded, did not produce any significant difference between PVT performance metrics when verified against the Control.

There was no significant difference in mean RT or mean 1/RT between the Melodic stimuli and the Control, however the Melodic stimuli did significantly reduced Lapses, False Starts, and produced a significantly improved PVT Performance Score than the Control. These results demonstrate that participants’ sustained attention has been significantly improved in the Melodic condition post-awakening. This suggests that alarm tones (not to fast nor to slow; 105 BPM) with melodically rhythmic features may be more successful in reducing the deficits experienced from *SI* post-awakening than counterparts devoid of melodic content (i.e. The Control stimulus we have tested).

The Rhythmic alarm tone did not produce any significant difference between PVT performance metrics compared to the Control indicating that a saliently rhythmic composition devoid of melody may be equally ineffective as a monotonous tonal beat sounded at 105 BPM concerning *SI* reduction.

We failed to reject the null hypothesis of equality between the subjective sleep attributes gathered for Test Group A (KSS, Hours Slept, Sleep Quality) in both test sessions (See Test Group A Results). Data thus reveals that during each test session there is no global effect of chronic sleep deprivation with respect participant performance. Acute sleep deprivation is unlikely considering the variability of individual differences in recommended sleep requirements as measured by the subject’s Typical Hours Slept per night, and Hours Slept prior to each test session (84, 85). For example, the mean percentages of all participants show that 50% report to maintain sleep epochs consistent with current recommendations (7+ hours per night), and 40% regularly sleep within or marginally below the ‘May be appropriate’ range (5 – 7 hours) (84). Prior to each test session 65% of participants slept in the recommended range (7+ hours) and 95% of all subjects sleep ‘May be appropriate’ (84). Additionally, the subjective Sleep Quality ratings show all participants (100%) report to have had at least an ‘Average’ to ‘Very Good’ night’s sleep.

The KSS reports do provide evidence of mild *SI* upon awakening in both test sessions (70% do not require any effort to stay awake upon arousal). This would be inconsistent with our PVT data if the Melody condition were assumed to be immediately effective upon awakening compared to the Control. However, due to the mild state of *SI* observed, the KSS may not be as sensitive to the effects auditory stimuli may elicit on vigilance comparative to the PVT. For example, Kaida and Abe (86) reported that when testing alert participants during a monotonous task while exposed to an ‘own name’ auditory condition, PVT Lapses were significantly improved relative to the other test conditions (including silent control), though there was no significant results observed in the KSS ratings to reflect the PVT data (86).

Test Group B’s (Rhythm, Control) self-reported sleep data indicates that it is unlikely participants in each test session exhibit chronic or acute sleep deprivation. For example, all participants in this group maintain sleep epochs that may be appropriate or marginally below recommendations (84). Additionally, 95% of participant sleep bouts preceding each test session may be suitable (84), with a majority (70%) reporting adequate sleep (7+ hours). The KSS and Sleep Quality measures indicate that the majority of participants were sleepy upon arousal and experienced an Average to Good night’s sleep, indicating that the effects of perceived *SI* may be mild upon awakening.

The results obtained from this study present new evidence for the potential of auditory alarms to effectively counteract the inhibiting symptoms of *SI*. Specifically, this study’s results support previous research showing a significant relationship between the reported melodicity of participants’ waking sound and a measure of perceived *SI* reduction (34).

One hypothesis we present for the lack of significant mean differences between the Melody and Control mean RT and mean 1/RT may be attributed to the beat intervals and perceived pace of both stimuli. In this study, the meter, beat, and temp of each stimuli are identical. We hypothesize that by retaining the existing tempo and increasing the beat intervals of the Control treatment within the Melody stimulus may produce a rise in the perceived ‘pace’ of the treatment, and in turn effect response time in a similar manner that faster tempo music has been shown to reduce RT (87–89) when compared to slower music. We extend this hypothesis to the Rhythm and Control results in Test Group B also.

Reductions in mean Lapses, False Starts and Performance score between the Control and Melody treatments in this study are significant, and may be attributed to the disparity in melodic content between the two stimuli. The Control stimulus was designed specifically as an unmelodic counterpart to the Melodic stimuli. This was achieved by retaining the Control as the beat of the phrase, and strategically composing a melodic contour around the Control. Research has shown that musical stimuli with melodic features (instrumental, Vivaldi’s Four Seasons ‘Spring’ concerto) increases task performance when compared to silent conditions (41, 44), and it is posited by Riby (44) that this may be a consequence of music’s ability to increase arousal and enhance cognition. Additionally, the mode in which the music is sounded has been attributed to improvements in performance (45), and that pitches in the range of the female human voice (~ < 2500 Hz) may be more successful in arousing sleeping humans than male (90). The frequency range of the melodic treatment in this study resides between 1318.5 Hz and 2637 Hz.

The contrast between the Control and Melodic stimuli in this study are consistent with these findings and may therefore have contributed to the improved vigilance of participants post awakening. Similarly, this examination may clarify the lack of significance between the Control and Rhythm stimuli. The Rhythmic treatment was composed to explore the effects of rhythm on *SI* in isolation void of a melodic contour. In this regard, the absence of melody may account for the insignificant results against the Control.

Future investigations into auditory countermeasures for *SI* may include the effects of tempo and melody, mode, or extend to additional auditory types including noise and the human voice. With respect to ecological inquiries the development of new methods and tools would assist in clarifying participant sleep profiles more efficiently, and would be advantageous in studies where controlled laboratory conditions are unsuitable or unattainable (e.g. The International Space Station).

This research presents evidence demonstrating that a musically melodically rhythmic alarm tone improves vigilance immediately upon awakening from a typical night’s sleep when compared to a metronomic alarm devoid of melody and with a restrained rhythmic contour. Additionally, this study’s results support previous research by McFarlane et al (34) showing a significant relationship between the reported melodicity of a participants’ waking sound and a measure of perceived *SI* (34). Taken together, these results highlight the potential importance of musical melody in waking sound design as an agent to counteract *SI*, and more broadly emphasizes the requirement for research which tests musical elements of stimulus design beyond the broader, more general music classifications such as genre.

Auditory alarms are a popular tool for assisted awakening. Our research provides evidence that may be utilized to produce effective auditory designs to counteract the unfavourable effects of *SI* for improved day-to-day awakening. In this study we thus provide the material required to precisely synthesize the stimuli we have employed, enabling future methodologies to further explore melody’s effect on *SI*.

## Ethics statement

All research methods, participant numbers (*N* = 20) considered appropriate for the study, and data collection were approved by RMIT’s University College Human Ethics Advisory Network (CHEAN) (ref: CHEAN B 21753-10/18). Informed consent was given online by all participants by way of participating in the study.

## Data availability, DOI

10.6084/m9.figshare.11905683

## Supporting Information

**S1. Table. Online questionnaire for each test session**.

